# Epistasis and physico-chemical constraints contribute to spatial clustering of amino acid substitutions in protein evolution

**DOI:** 10.1101/2020.08.05.237594

**Authors:** Andrew M. Taverner, Logan J. Blaine, Peter Andolfatto

**Affiliations:** Lewis-Sigler Institute for Integrative Genomics, Princeton University, Princeton, NJ 08544; Department of Molecular Biology, Princeton University, Princeton, NJ 08544; Department of Biological Sciences, Columbia University, New York, NY 10027

## Abstract

The causes of rate variation among sites within proteins are as yet poorly understood. Here, we compare the spatial autocorrelation of non-synonymous substitutions among species within diverse phylogenetic groups: *Saccharomyces, Drosophila, Arabidopsis*, and primates. Across these taxa, we find that amino acid substitutions exhibit excess clustering that extends over a 20-30 codon length (10-20 Angstrom distance) scale. We show that these substitutions cluster more strongly and exhibit compensatory dynamics within species lineages but exhibit patterns of convergent evolution between lineages. We evaluate a simple model of thermodynamic constraints on protein folding and conclude that it is unable to recapitulate the observed spatial clustering of substitutions. While pairs of substitutions with the strongest epistasis tend to spatially cluster in these simulations, the magnitude and length scale are smaller than that observed in real data. Additionally, we show that the pattern of convergent substitution is also not expected under this model, suggesting it is likely caused by factors other than these simple thermodynamic constraints. Our results support a prevalent role for epistasis and convergent evolution in shaping protein evolution across the tree of life.

## Introduction

Evolutionary rate variation is closely linked to protein structure and function, and understanding its causes is essential to understanding the mechanistic basis and functional relevance of protein evolution [1]. For example, several algorithms used to identify pathogenic substitutions in humans employ evolutionary rate variation information [2,3]. However, as most of these methods assume a simple model of stabilizing selection and independence among sites, a more detailed understanding of the factors contributing to rate variation would likely increase the sensitivity and specificity of such approaches [1]. In addition, incorrect assumptions about how evolutionary rate varies within and among proteins can lead to incorrect inferences of phylogenetic relationships and evolutionary rates [4] and the mode and intensity of selection acting on proteins [5–7].

Despite its importance, the causes of evolutionary rate variation among proteins and among sites within proteins are currently not well understood [8]. A number of studies have examined correlations between various protein properties and differences in evolutionary rate variation [9–11]. These studies have identified structural properties such as relative solvent accessibility, weighted contact number, and protein flexibility as major determinants of evolutionary rate [8]. Broadly speaking, the interiors of proteins, with higher levels of contact with other residues and more physical constraints, exhibit higher levels of evolutionary constraint. Beyond their structural integrity, proteins are additionally constrained in terms of their functional requirements. Not only is the active site of proteins highly constrained, constraint decreases roughly linearly with distance from the active site [12,13]. Despite studies on variation in constraint, there has been a relative dearth of studies investigating the effect of interactions among amino acid residues on evolutionary rate variation within proteins. While properties such as relative solvent accessibility and weighted contact number depend on interactions between amino acid residues, it is surprising that the direct effect of interactions between amino acids on evolutionary rate variation has not been more studied.

Emerging evidence suggests that genetic interactions among amino acid residues within a protein, called “intramolecular epistasis”, are a strong determinant of patterns of protein evolution [14]. Epistasis refers to a broad genetic phenomenon by which the phenotypic or fitness effects of given substitutions are dependent on the genetic background on which they arise [15]. Given that steric and allosteric physical interactions among amino-acid substitutions within a protein are common, it follows that biochemical and evolutionary fitness effects of amino acid substitutions can be sequence-context dependent [16,17]. This view of constraints on protein evolution was first formulated as the “covarion” model [18], which proposes that only a limited number of amino acid substitutions can be tolerated on the current background of a protein, and that each substitution changes the pool of subsequently tolerable substitutions.

Covarion-like epistasis models are relevant to understanding protein evolution in two contexts. Firstly for its implications for how Darwinian positive selection is expected to proceed in the evolution of proteins with new or enhanced functions [19–22]. However, similar epistatic constraints can also be relevant to neutral (or nearly-neutral) protein evolution [23–25]. Specifically, epistasis can limit protein evolution under either scenario by enforcing a limited set of viable substitutions, such that each substitution still results in a functional protein [22]. While primarily explored by simulation in the context of constraints on protein folding [24,25], covarion-like models have been shown to be consistent with molecular evolutionary patterns observed in a number of proteins, including bovine RNase, cytochrome C, and Cu, Zn superoxide dismutase [18,26,27] and supported by functional experiments on proteins involved in antibiotic resistance [22], steroid-receptor ligand specificity [28], viral immune escape [19] and toxin-insensitivity [29].

Since direct contacts between amino acid residues are the most obvious source of interactions, and primary sequence is a strong predictor of physical proximity in threedimensional protein structures [30], one might expect that epistatic effects among amino acid substitutions could contribute to local spatial evolutionary rate variation. Intriguingly, the analysis of small datasets of compensatory substitutions in proteins revealed that they tend to spatially cluster within proteins [31,32].

Two studies that have explored the spatial autocorrelation of non-synonymous substitutions using genome-scale data and came to remarkably similar conclusions [30,33]. Specifically, both studies confirmed that non-synonymous substitutions cluster spatially within proteins and that the degree of constraint in the protein is positively correlated with the strength of clustering [34]. Callahan et al. [30] further used closely-related *Drosophila* species to show that clustering is more pronounced for pairs of amino-acid substitutions that occur in the same species lineage and that there is an enrichment of charge-compensating substitution pairs on a length scale of ~20 codons, strongly implicating epistasis as a contributing factor. Intriguingly, the analysis of Callahan et al. [30] also found an enrichment of charge-reinforcing substitution pairs occurring in different lineages over a similar length scale, suggesting convergent evolutionary dynamics may also contribute to clustering.

Studies of genomic patterns of amino acid substitution clustering have thus far been carried out in *Drosophila* which are obligately sexual species with large population sizes that exhibit strong signatures of adaptive protein evolution [35]. However, such patterns are not observed as strongly in other taxa. To evaluate the generality of patterns of clustered amino acid substitutions and their link to compensatory, convergent, and adaptive evolution, we investigate clustering patterns in a diverse set of four taxonomic groups, comprising *Saccharomyces, Drosophila, Arabidopsis*, and primates, which vary broadly in mating system, population size, and other population genetic parameters. We also carry out simulations of constraint on protein folding to ask whether such constraints may be sufficient to explain observed patterns of amino acid clustering.

## Results

Using multisequence alignments of *Saccharomyces*, *Drosophila*, primates, and *Arabidopsis*, we evaluate whether the patterns of spatially clustered non-synonymous substitutions observed in Drosophila are generalizable to other diverse taxa. To quantify spatial clustering, we measure the observed number of pairs of: two non-synonymous substitutions (DNDN) and two synonymous substitutions (DSDS) at each distance, *x*, in codons, compared to the null distribution of randomly distributed substitutions. Our clustering metric has an intuitive interpretation: it describes the increased likelihood of observing a second substitution of a certain type, conditioned on an initial mutation separated by *x* codons. Furthermore, we see an enrichment in this score, typically peaking at *x* = 1 and decreasing over approximately 20 codons. Following the convention of Callahan et al., we refer to this as “clustering” [30].

### Clustering of non-synonymous substitutions occurs in all four taxa

In all pairwise comparisons within the four taxa, we observe a peak in our clustering metric for DNDN at the adjacent amino acid with respect to the focal substitution. From here, the clustering rapidly decreases over roughly 20 codons. In fact, in both *Saccharomyces* and *Drosophila*, there is a nearly 1.7 times increased chance to find an adjacent non-synonymous substitution, conditional on a focal non-synonymous substitution, relative to the baseline rate of non-synonymous pairs 100 codons apart. This clustering in the *Drosophila* clade recapitulates the findings of Callahan et al. [30], and further extends the finding to *Saccharomyces*, primates, and *Arabidopsis*.

Notably, DSDS shows far less clustering compared to DNDN. In *Saccharomyces*, DSDS shows no clustering, while *Arabidopsis*, *Drosophila*, and primates show a slight clustering peak (Figure 1). This is largely consistent with a pattern of uniformly distributed synonymous substitutions. This indicates that the clustering observed is a phenomenon relevant to the protein sequence itself, given that only DNDN strongly shows this clustering pattern. These results are consistent with a model in which amino acid substitution pairs interact and experience selection at the protein level.

**Figure 1:**
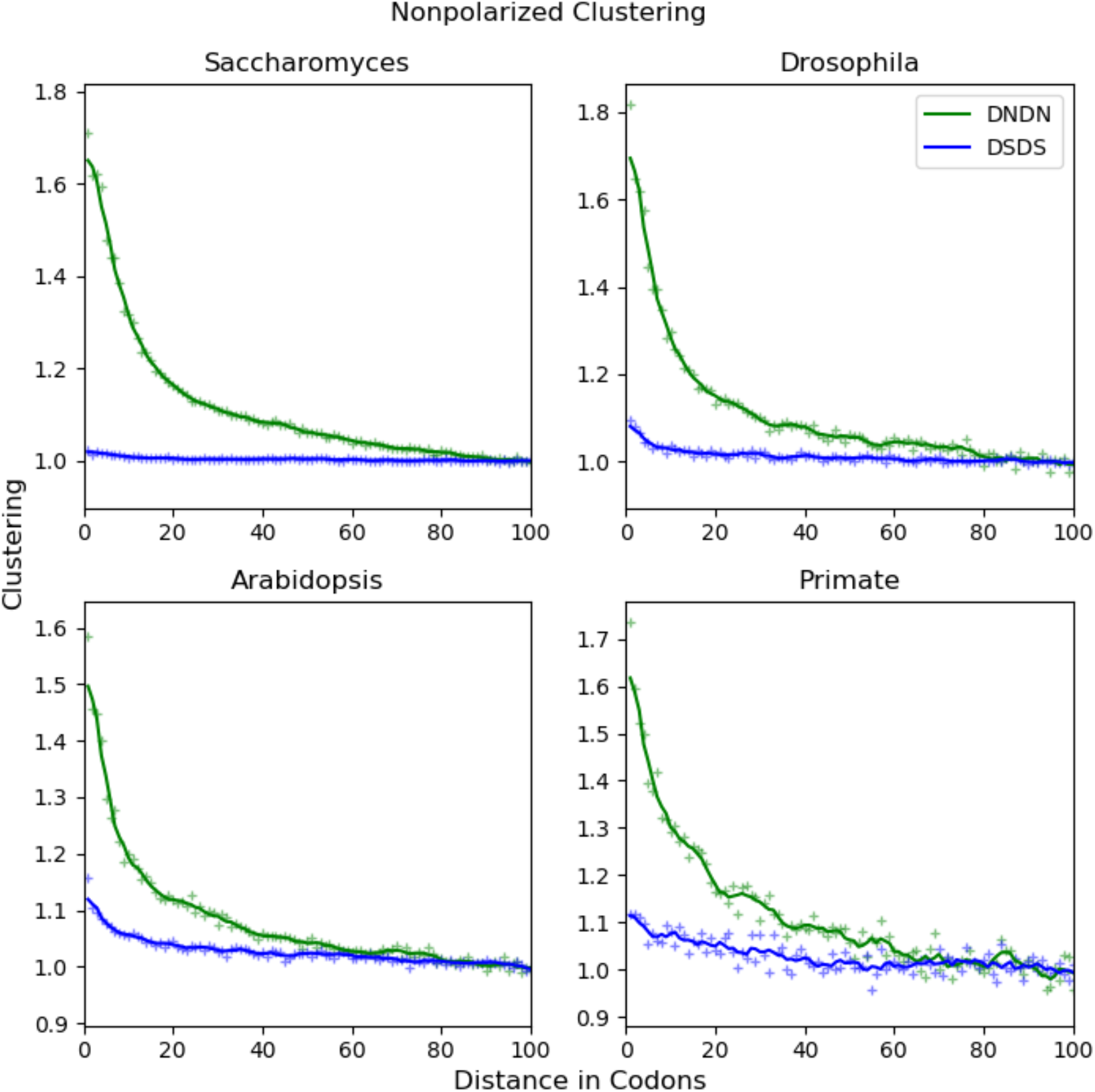
Non-synonymous substitutions cluster spatially within proteins compared to synonymous substitutions. Clustering of non-polarized substitution pairs: two non-synonymous (DNDN – green) and two synonymous (DSDS – blue). Non-synonymous substitutions show significant clustering compared to uniform expectation (*p* < 10^−300^, except for primates: *p* = 1.1 × 10^−l09^) and compared to the distribution of synonymous substitution clustering (*p* < 10^−300^, except for primates: *p* = 2.6 × 10^−78^). These distributions represent the increased likelihood of observing a second substitution, conditional on a focal one, *x* codons away, normalized to 1 at long-codon distance. In all four taxa, DNDN peaks at *x* = 1, indicating that the maximum chance of observing a second non-synonymous substitution, given a focal non-synonymous substitution, occurs at the adjacent amino acid. Furthermore, DNDN clustering rapidly decreases over a 20-30 codon length scale. DSDS serves as a control, and reassuringly, show little or no clustering. Raw data is indicated with a “+”; lines are smoothed using a 5-codon sliding window. Non-polarized clustering is shown for the following comparisons: *S. cerevisiae* and *S. mikatae; D. melanogaster* and *D. simulans; A. thaliana* and *A. lyrata; H. sapiens* and *P. pygmaeus abelii*.

By quantifying clustering at the level of primary sequence, we are assuming that distance along the primary sequence is a good proxy for 3D distance in folded protein structures [30]. To verify this, we calculated clustering using the atomic distance (in angstroms) between the alpha carbon in the amino acid residue. Protein structures were available for only 37% and 38% of proteins in our primate and *Saccharomyces* datasets, respectively. Despite this limitation, both *Saccharomyces* and primates (the only two taxa for which sufficient 3D data exists), non-synonymous substitution pairs cluster over 10-20 Å distance while synonymous pairs do not. Additionally, distance in the primary sequence correlated with distance in 3D space, particularly for substitutions on a length scale of 20 codons (Figure 2).

**Figure 2:**
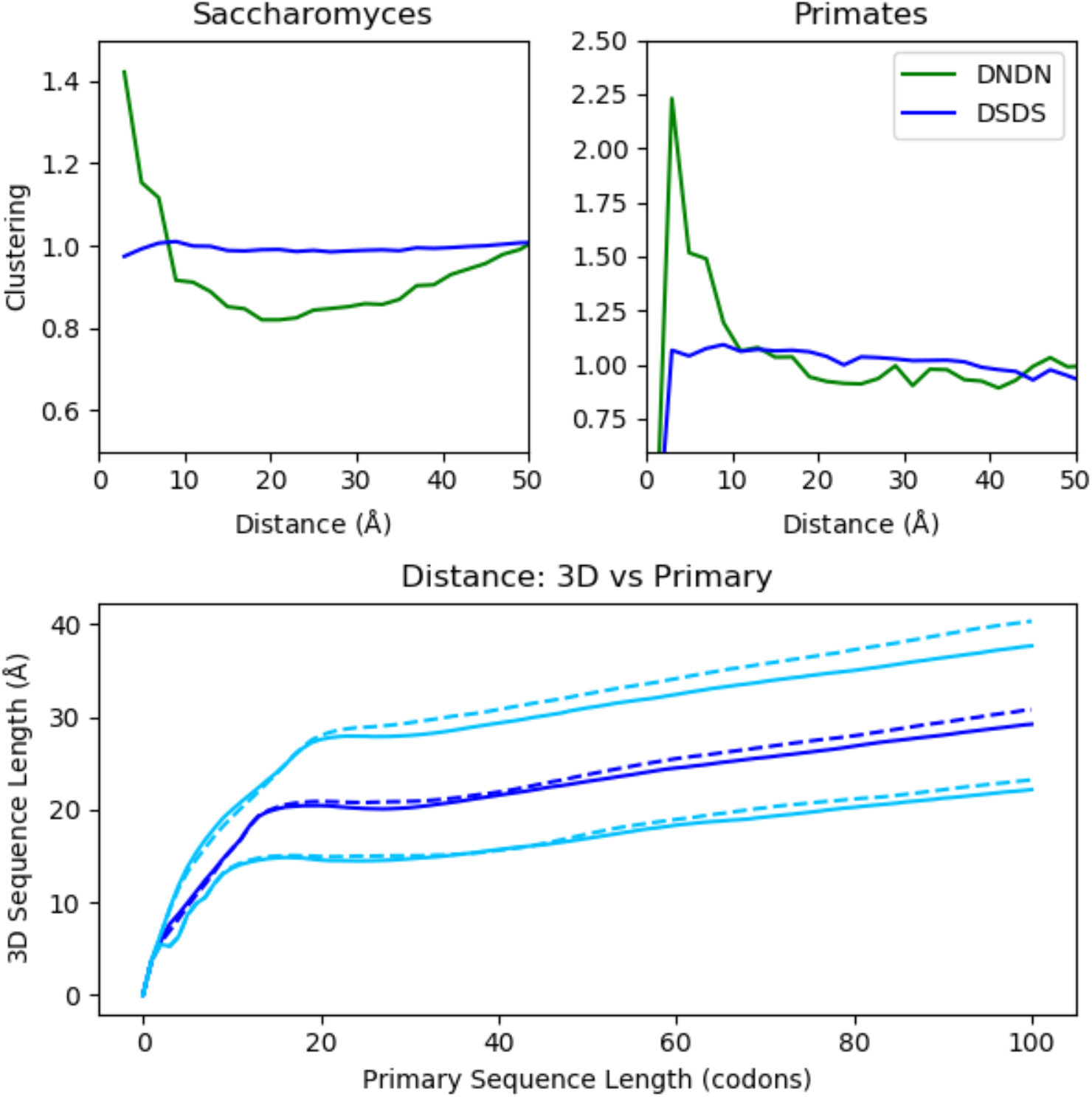
Top row: Using angstroms as the distance metric between amino acids (instead of using codons as a proxy) recapitulates the findings in Figure 1: non-synonymous substitutions tend to be clustered, while synonymous ones do not. 3D Clustering is shown for *Saccharomyces* (*S. cerevisiae* and *S. mikatae*) and primates (*H. sapiens* and *P. pygmaeus abelii*). Distances between amino acids are binned in 2Å increments. No mutations were observed in the primate data for the 0-2 Å bin, due to the rarity of substitutions at this small length scale and the lack of available data (i.e. the 3D structure of the protein was resolved and the gene alignment passed filtering critera). Bottom row: Distance in 3D space in Å vs distance in primary sequence in codons. The relationship is shown for *Saccharomyces* (dashed line) and primates (solid line). Median distance (dark blue) and 1^st^ and 3^rd^ quartiles (light blue) are shown. Together, this suggests that using distance between amino acids in codons is a good proxy for distance in 3D space.

### Non-synonymous substitutions cluster more strongly within a species lineage than between lineages

While we have shown (above) that species-distinguishing amino acid substitutions cluster in sequence and 3D structural space, such patterns could result from simple variation in evolutionary constraint related to protein domain structure and function. Thus, following [30], we next consider the spatial pattern of substitutions along species lineages. Substitutions were polarized into the species lineage in which they arose with an appropriate out group, using a parsimony-based method. We repeated the clustering analysis using only substitutions from one lineage to calculate within lineage clustering, while between lineage clustering was calculated using a focal substitution from one species and a second substitution from the other species. Amino acid clustering between lineages must be independent events so between lineage clustering can only be explained by a shared property, like reduced constraint, between homologous proteins. Thus, we would expect between lineage clustering to be roughly equal to within lineage clustering if these patterns were due solely to a reduction in constraint in certain regions of proteins.

In all four taxa, clustering is observed both in within lineage and between lineage clustering and as before, clustering peaks at *x* = 1, and decreases over the next 20 codons. In *Saccharomyces*, *Arabidopsis*, *Drosophila*, and primates, within lineage clustering is consistently greater than between lineage clustering. Additionally, in both *Saccharomyces* and *Drosophila*, within-species clustering excess appears to be approximately equal. Interestingly, *P. pygmaeus abelii* show drastically higher amounts of within lineage clustering, even compared *H. sapiens*, though these results are relatively noisy (Figure 3). This pattern of elevated within-lineage clustering is consistent with a model in which reduced constraint in certain regions of proteins is responsible for some, but not all, of the clustering pattern we observe.

**Figure 3:**
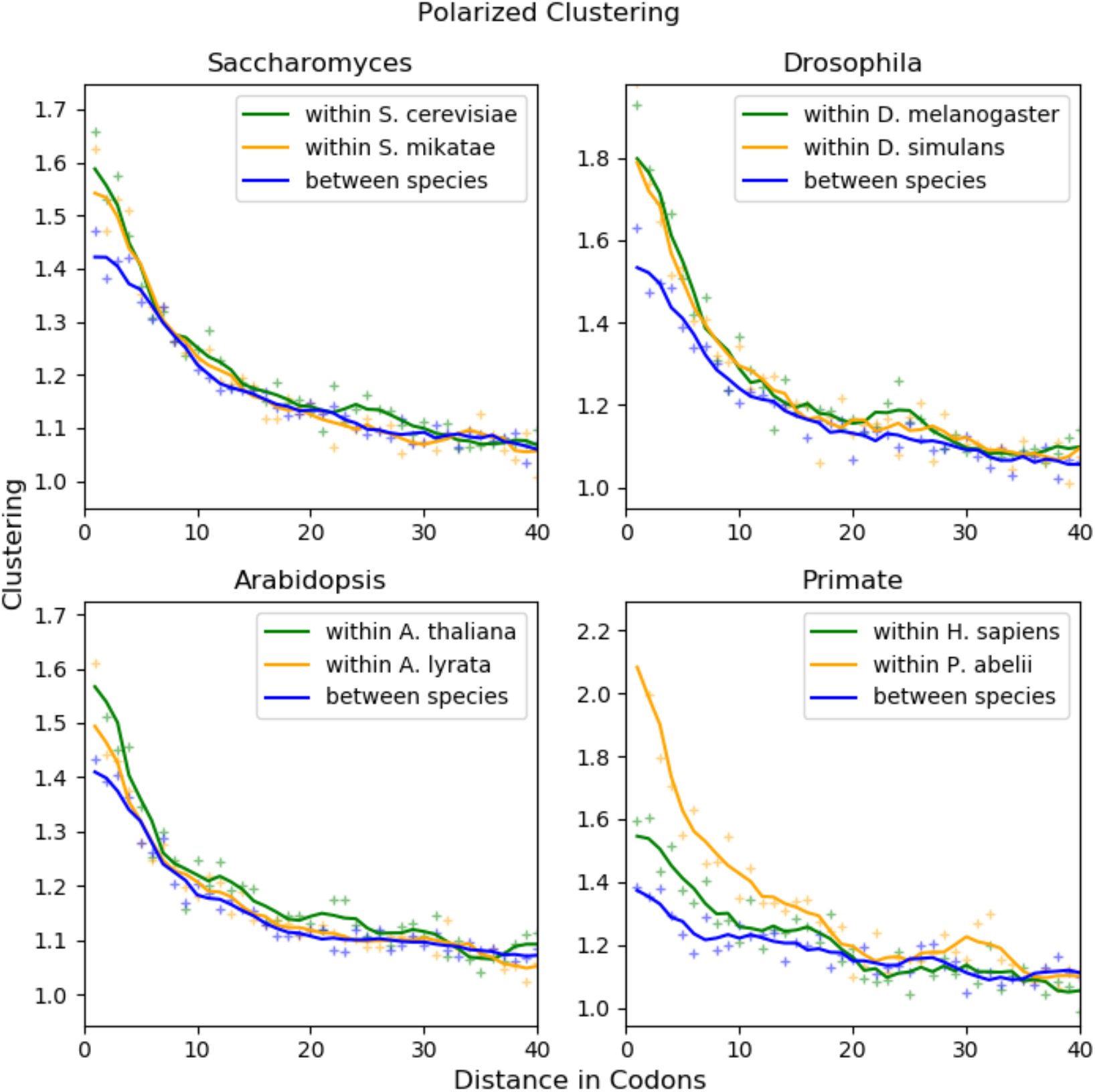
Pairs of non-synonymous are more clustered when they both occur within a species than when they occur between species. Pairs of non-synonymous substitutions occurring within in the same species shown in green and orange. Pairs occurring between species are shown in blue. In all four taxa, within species clustering is significantly greater than between species clustering, indicating an excess of within species clustering (*p* < 10^−6^, except for within *H. sapiens* clustering, for which *p* = 0.01. See Table S2 for exact values). Calculating between species clustering breaks any interactions the substitutions might have on one another, thus elevated within-species clustering is consistent with a model in which substitutions interact with one another. Again, raw data is shown with a lines are smoothed using a 5-codon sliding window. Substitutions are polarized into the lineage in which they arose using parsimony and an appropriate outgroup: *S. cerevisiae* and *S. mikatae*, outgroup: *S. kudriavzevii*; *D. melanogaster* and *D. simulans*, outgroup: *D. yakuba*; *A. thaliana* and *A. lyrata*, outgroup: *C. rubella*; *H. sapiens* and *P. pygmaeus abelii*, outgroup: *P. anubi*

### Evidence for compensatory charge changes within lineages and convergent evolution

We considered properties such as amino acid charge, polarity, and size and looked for consistent, repeated patterns of clustering among substitutions causing changes of these properties. For example, if one amino acid substitution is positive with respect to the ancestral state, another nearby amino acid may become negative with respect to the ancestral state to compensate for this change. Alternatively, if this second amino acid substitution is positive with respect to the ancestral state, this pair of amino acids would reinforce the charge change. We term these two possibilities as compensatory and reinforcing pairs of substitutions, respectively. To test this, the fraction of polarized, non-synonymous substitutions pairs, resulting in either compensatory or reinforcing charge changes were analyzed.

In three of the four taxa (*Saccharomyces, Drosophila*, and *Arabidopsis*), there is a clear pattern of enrichment of charge compensating pairs, over a length scale of 5-15 codons. This clustering, however, only occurs within lineage, while there appears to be no clustering between lineages. Furthermore, in these three taxa, roughly 7-8% of non-synonymous pairs are charge compensating, compared to the observed baseline of approximately 5% that appears to be largely consistent at greater length scales (>20 codons). This suggests that maintenance of local charge within a protein is an important factor affecting protein evolution.

In contrast, when looking at pairs of substitutions with reinforcing charge changes, the opposite pattern appears. No obvious reinforcing clustering can be observed within lineage, but reinforcing clustering is observed between lineage. Again, this pattern is found in *Saccharomyces, Drosophila*, and *Arabidopsis*, but not in primates. Similar to compensatory clustering, reinforcing clustering peaks at *x* = 1, and is observed over 5-15 codons (Figure 4). This reinforcing clustering indicates that similar molecular changes are occurring in similar regions in homologous proteins, suggesting that molecular convergent evolution may be widespread or alternatively that certain regions within proteins are tolerant to certain types of substitutions. We observe similar, though less obvious, patterns for both size and polarity reinforcing and compensatory clustering in *Saccharomyces, Arabidopsis*, and *Drosophila*. However, this within lineage, size compensating clustering appears to be mostly absent in *D. melanogaster* (Supplementary Figure 1 & 2).

**Figure 4:**
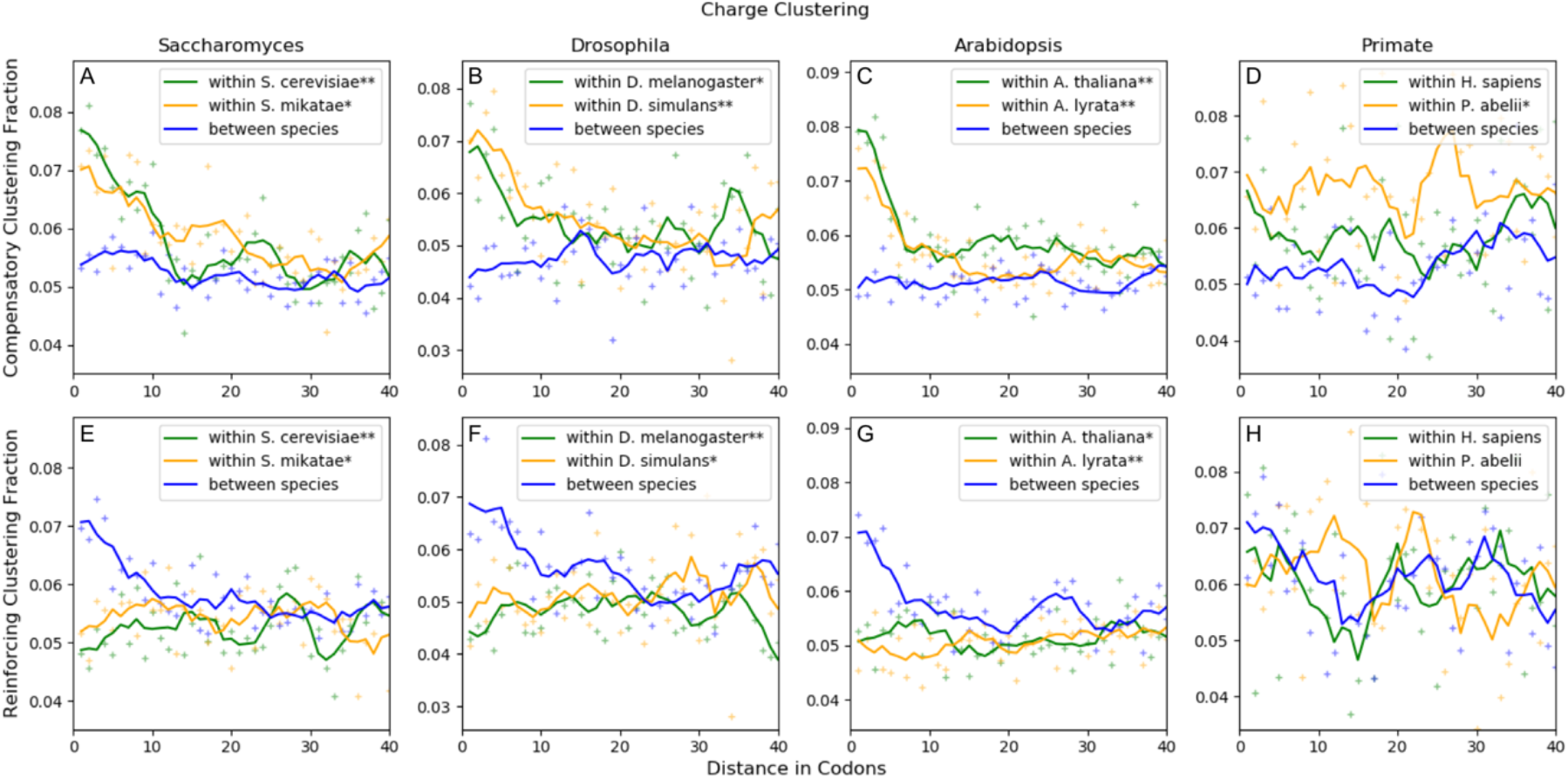
Pairs of non-synonymous substitutions causing compensatory charge changes are clustered within species. In contrast, pairs of non-synonymous substitutions causing reinforcing charge changes are clustered between species. The top row of panels show the fraction of charge compensating pairs of non-synonymous substitutions (A, B, C, D), while the bottom row show the fraction of charge reinforcing pairs of non-synonymous substitutions (E, F, G, H). Again, pairs occurring within species are shown in green and orange, while pairs occurring between species are shown in blue. Compensatory charge changes tend to be significantly clustered within a species lineage, but not between lineages. In contrast, reinforcing changes are tend to be significantly clustered between species but not within species. P-values are indicated in the legend by: ** *p* < 10^−10^; * 10^−10^ < p < 0.05. (See Table S3 for p-values). The following comparisons are shown: A, E) *S. cerevisiae* and *S. mikatae*, outgroup: *S. kudriavzevii*. B, F) *D. melanogaster* and *D. simulans*, outgroup: *D. yakuba*. C, G) *A. thaliana* and *A. lyrata*, outgroup: *C. rubella*. D, H) *H. sapiens* and *P. pygmaeus abelii*, outgroup: *P. anubis*.

### Simulations reveal that selection to maintain folding stability is unable to quantitatively recapitulate clustering

Given that we observe widespread compensatory clustering (mostly for charge), we wanted to disentangle whether these changes were due to purifying selection working to maintain the protein function or positive selection. To this end, we used FoldX [36] to calculate the change in folding energy of simulated substitutions in three proteins as a proxy for fitness. We chose the lysine-, arginine-, ornithine-binding protein (1LAF), NmrA-like family domain containing protein 1 (2WM3), and mini spindles TOG3 (4Y5J) for our simulations (PDB ID is given in parentheses). We shall refer to these proteins by their IDB ID from this point forward for brevity.

We first assessed whether FoldX was able to effectively simulate epistasis caused by selection by simulating pairs of substitutions, selecting for wild-type folding energy. We compared the ΔΔG_1_ + ΔΔG_2_ against the ΔΔG_1,2_, that is, the change in folding energy between the sum of the first and second substitutions alone against the change in folding energy of the double mutant. Under a purely additive model, we would expect ΔΔG_1_ + ΔΔG_2_ to be equal to ΔΔG_1,2_. When we simulate selection toward wild type folding stability, the correlation coefficient between the additive change in folding energy of the single mutants vs the double mutant is r^2^ = 0.62 with a slope of 0.79, indicating some level of non-additivity, or epistasis between substitutions (Figure 5). This pattern holds for each of the proteins individually so this pattern is not being driven by any one protein (Supplementary Figure 5). We also examined the distribution of substitutions across the proteins, and found a non-uniform distribution of substitutions, as expected, given that different sites should have different levels of constraint (Supplementary Figure 4).

**Figure 5:**
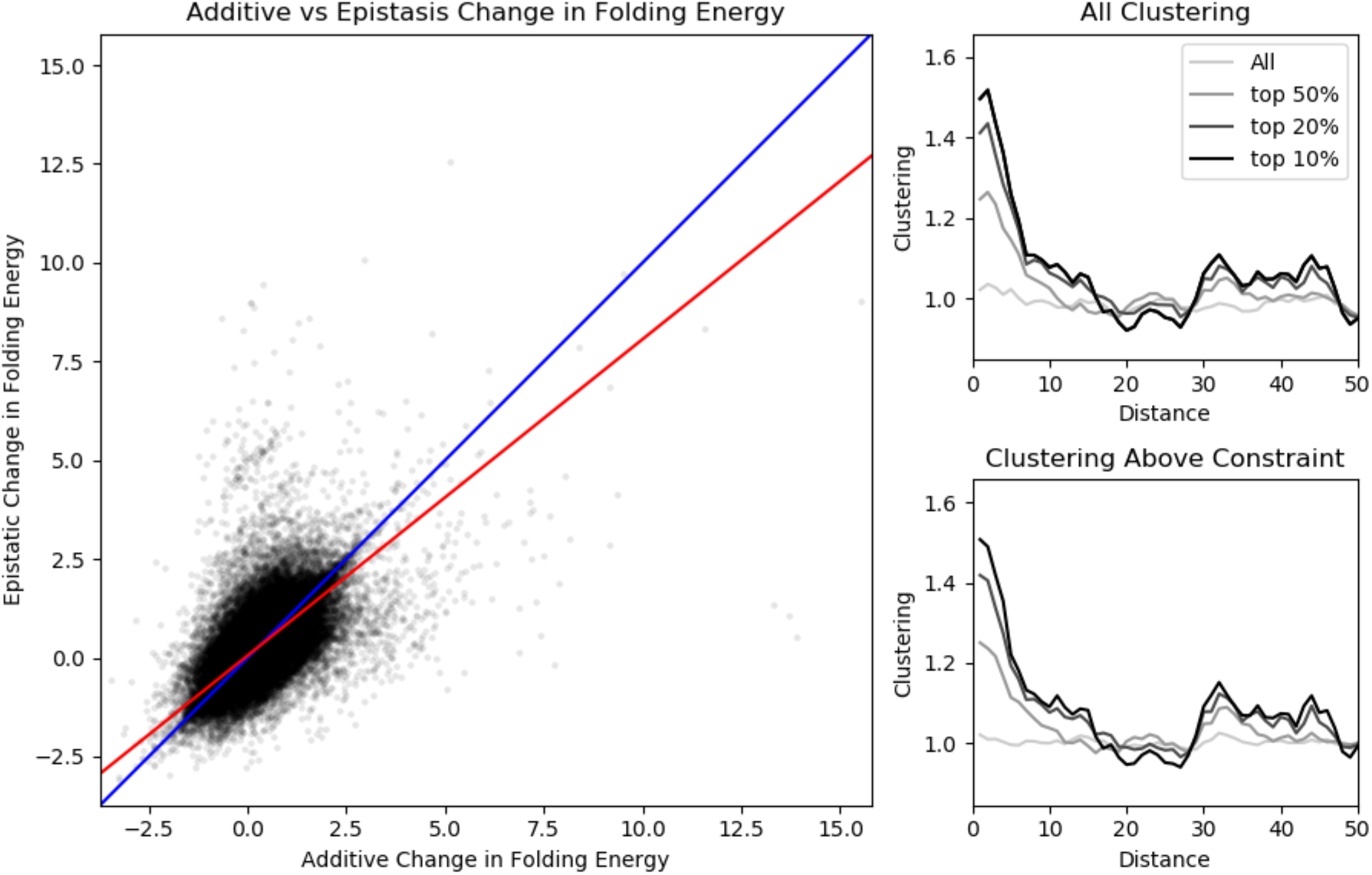
A) Epistasis between pairs of simulated amino acid mutations is apparent when selecting to maintain the wild-type folding energy. Each point represents one pair of simulated mutations in one of the three proteins. The additive change in folding energy (ΔΔG_1_ + ΔΔG_2_, x-axis) is on average higher than the epistatic change in folding energy (ΔΔG_1,2_, y-axis). The 1:1 line is shown in blue, while the red line shows the linear regression of the points. Without interactions between amino acid mutations, the additive and epistatic change in folding energy is expected to be equal. However, the epistatic change in folding energy is less than the additive expectation. B,C) Clustering between all pairs of simulated mutations show no apparent clustering. However, after enriching for epistatic pairs of simulated mutations, clustering is observed, even taking only the top 50% of the most epistatic pairs of mutations. Panel B shows clustering due to any causes, while panel C shows clustering after normalizing for variation in constraint across the proteins. Panels B and C look qualitatively similar, so variation in constraint is not a significant driver of the simulated clustering. In all panels, simulations across all three simulated proteins (1LAF, 2WM3, and 4Y5J) have been aggregated.

Given that FoldX was able to simulate epistasis, we conducted simulations of pairs of substitutions, again in a stepwise fashion, selecting to maintain wild type folding energy. We reasoned that FoldX would be able to more accurately calculate the change in folding energy given fewer simulated substitutions. This had the additional benefit of speeding our simulations so we could gather a larger sample. Our clustering metric was centered around 1, and we did not observe clustering either overall or when accounting for clustering that might be due to variation in protein constraint (Figure 5).

Lastly, we wanted to know whether epistasis was sufficient to cause clustering; or in other words, would a subset of substitutions that display epistatic interaction be clustered spatially. We filtered for the most epistatic pairs of substitutions; ones with a non-additive ΔΔG (|ΔΔG_1_ + ΔΔG_2_ - ΔΔG_1,2_| >> 0). In the subset containing the 90^th^ percentile of the most epistatic pairs, we found clustering patterns that were similar to the ones we observed in the real data: the distribution peaked at the adjacent amino acid at approximately 1.5 and decreased over the next 10-15 amino acids (Figure 5). This indicated approximately 50% more substitutions at the adjacent amino acid than would be expected due solely to selective constraint. Furthermore, we observed that filtering out epistatic substitution pairs disproportionately removed pairs that were “far apart”, more than ~15 codons away (Supplementary Figure 8).

## Discussion

In this study, we show that amino acid substitutions tend to alter the local evolutionary rate across a protein, leading to an accumulation of proximate amino acid substitutions. This clustering peaks at the adjacent amino acid and decreases over a span of approximately 20-30 codons. As we expect selection acting on amino acid substitutions is driving the clustering we observe, the fact that we observe very little clustering of and DSDS is reassuring. However, clustering could also be due to hitchhiking of nearly-neutral substitutions along with a positively selected substitution, multi-nucleotide mutation (MNM) events, or decreased constraint in regions of proteins. This is unlikely, however, as MNM are expected to occur over quite short length scales [37]. Furthermore, we can eliminate hitchhiking as a driver of the clustering pattern we observe, as this would affect synonymous substitutions as well. We ultimately conclude that a non-negligible amount of the clustering we observe is driven by selection.

Another possible cause of this clustering could be variation in constraint among functionally coherent domains of proteins. For example, the active site of proteins tends to be highly conserved, while amino acid residues on the exterior of globular proteins are less constrained [9–13]. If this were the case, we would expect substitutions to cluster in the same unconstrained regions across our multi-species alignments. When calculating clustering across lineages, we break any interactions that may occur between amino acid substitutions. We find that amino acid substitutions cluster more strongly within lineage than between lineage, showing that clustering does not occur solely as a result of an accumulation of substitutions within unconstrained regions of proteins. Thus, we expect the excess within-lineage clustering to be due to interactions between amino acid residues, driven by selection on these epistatic interactions.

Previous studies support these findings as they have noted similar patterns of substitutions co-occurring spatially more frequently than expected if the substitutions had occurred randomly. Two studies found that in HIV evolution and mouse-rat divergence, even within a single codon, double and triple substitutions occur at a higher rate than expected [38,39]. Other studies have also found longer-range signals of co-occurring substitutions. For example, two studies found that non-synonymous substitutions auto-correlate spatially along the primary sequence of proteins, while synonymous ones do not [30,33]. It has been shown that such compensatory substitutions cluster both with one another and are also nearby the deleterious substitution, which could be one explanation of the clustering pattern we observe [31].

One key assumption we make is that distance along the primary sequence is a good proxy for distance in 3D space. While a high correlation exists between distance in primary sequence with distance in 3D space at short length scales (Figure 2, and see [30]), as an additional sanity check, we calculated 3D clustering in *Saccharomyces* and primates (Figure 2). Indeed, we found that substitutions tend to cluster over ~10Å distances, confirming that clustering occurs between proximate residues and that primary sequence distance is a reasonable proxy for 3D distance.

Next, we chose to examine whether there are any trends with respect to the amino acid property changes and found that there are a greater than expected number of charge compensating substitution pairs within a lineage that occur in a 5-15 codon window. Previous studies have observed charge compensation in evolution that are consistent with this finding. In influenza evolution, an arginine residue and glutamic acid residue (positively and negatively charged, respectively), are both substituted with glycine, a neutrally charged amino acid [19]. These amino acids are separated by 9 residues and these changes would maintain the local charge of the protein. Similarly, in rat RNase A, adjacent amino acid residues glycine and serine are, in bovine RNase A, aspartate and arginine, respectively [26]. Fitch and Markowitz speculate that this transition is only possible because the negative charge of aspartate and positive charge of arginine neutralize one another, preventing from disrupting other residues that are important for protein function. In addition to the compensatory charge clustering that is consistent with the literature, we find that charge reinforcing substitution pairs are overrepresented between lineages. This seems to indicate that similar types of substitutions are occurring between the two species, which could be an indication of parallel or convergent evolution. Alternatively, it could indicate that regions of proteins are only tolerant to certain types of substitutions, but this would also be evidence of epistatic interactions between these substitutions and the entire protein.

Similarly, we found size compensating clustering in three of the four taxa we examined. This is also consistent with previous findings: Fitch and Markowitz [26] observed in rat RNase A, an isoleucine and leucine, while bovine RNase A had a valine and methionine. In this example, the volume of the pair of amino acid residues is maintained, as valine is a smaller amino acid than isoleucine, while methionine is larger than leucine. However, we find that the patterns of compensatory and reinforcing clustering for size and polarity are less apparent than for charge. This may indicate that amino acid charge is more important than size and polarity in influencing protein evolution.

Previous studies have shown that maintaining the stability of proteins through epistasis can be an important factor in evolution. For example, a substitution that increases the stability of a protein may not have much of a fitness effect, as the protein is already sufficiently stable to function, but it may rescue the effect of a deleterious, destabilizing mutation [19]. Despite this expectation, we were unable to recapitulate any clustering signal in our simulations of protein folding stability using FoldX. While we do find significant clustering once we subset pairs of substitutions that interact epistatically, the magnitude and length scale of this clustering are smaller than observed in real data. This seems to indicate that, while purifying selection to maintain the folding-stability of the protein via epistasis does cause substitutions to spatially auto-correlate, it seems unlikely to be a significant contributor to the within-species compensatory dynamics and between species convergence among proximate substitution documented here.

On the whole, this may not be surprising given a few important caveats of our simulations. The most notable among these is that, given limits on our ability to computationally model folding-stability, we carried out simulations for three short proteins that may not be particularly representative. In fact, the proteins we selected for simulation are shorter than any of those we used to calculate clustering. Also, while thermodynamic constrains are important to protein function (for example see [19]), it is possible that they are a poor proxy for fitness, as there are other factors that contribute to protein function. Simulations of folding-stability do not account for critically important constraints near ligand-binding sites and or sites important for protein-protein interactions, as would be important for modelling, for example, hemoglobin [40,41].

An interesting alternative explanation for the patterns of within-lineage spatial clustering of substitutions is that they may reflect dynamics associated with adaptive evolution. For example, adaptive amino acid substitutions for resistance to cardiac-glycoside toxins have often been documented at ATPa (a subunit of Na+,K+-ATPase) in variety of insect species that feed on plants that produce these compounds [42,43]. In many cases, this involves repeated occurrences of three amino acid-substitutions that are within 11 residues of each other [29]. While two of these substitutions were previously known to confer cardiac glycoside-resistance to the protein (reviewed in [42]), both normal enzyme activity and overall animal fitness depend on a “permissive” substitution at the third site. Similar interactions among proximate substitutions in an adaptive context have been observed for the nicotinic acetylcholine receptor resistance to epibatidine in Neotropical poison frogs and high-altitude adaptation of hemoglobin in birds as well as pikas [40,41,44]. The frequency of adaptive protein evolution in Drosophila [35,45] suggests that this scenario is plausible, however it’s importance in yeast [46], Arabidopsis [47], and humans [48] is less clear. Distinguishing between adaptive and non-adaptive causes for within-lineage amino acid substitution clustering will be an interesting topic for future investigations.

## Methods

### Alignments

We used multisequence, coding sequence (CDS) alignments from four different taxonomic groups in this analysis: *Saccharomyces, Drosophila, Arabidopsis*, and primates. The *Saccharomyces* dataset consisted of five species: *S. cerevisiae, S. paradoxus, S. kudriavzevii, S. mikatae*, and *S. bayanus* from http://www.saccharomycessensustricto.org/cgi-bin/s3.cgi?data=Orthologs&version=current[49]. After removing CDS alignments lacking a *Saccharomyces* Genome Database (SGD) name and removing CDS alignments with duplicated SGD names, 5,155 genes remained 4,291 (83%) of which passed our filtering criteria and were used in the clustering analysis. From the *Saccharomyces* dataset, we quantified clustering in *S. cerevisiae* vs *S. mikatae* using *S. kudriavzevii* as the outgroup. The *Arabidopsis* dataset consisted of 16,263 CDS alignments for three species. Of these 16,263 genes, 13,234 (81%) passed our filtering criteria and were used in the clustering analysis. We quantified clustering in *A. lyrata* and *A. thaliana* using *C. rubella* as the outgroup [50]. The primate data consists of a subset of the 20 species alignment from the UCSC genome browser (UCSC Genome Browser: https://hgdownload.soe.ucsc.edu/goldenPath/hg38/multiz20way/). This dataset consistent of 37,372 transcripts and after selecting the longest transcript that also passed our filtering criteria, 9,516 genes remained. We quantified clustering in Human (*Homo sapiens*) and Orangutan (*Pongo pygmaeus abelii*) using Gibbon (*Nomascus leucogenys*) as the outgroup. Clustering analyses were performed in Python (code available at: https://github.com/andrewtaverner/clustering), using Biopython to parse the multisequence CDS alignments [51].

The *Drosophila* dataset was the only dataset for which a multispecies CDS alignment did not exist so we created a multispecies alignment for this taxa from reference genomes. The *Drosophila* dataset consisted of three species: *D. melanogaster*, version 6.11 from FlyBase [52] and *de novo* assemblies of *D. simulans* w501 and *D. yakuba* Tai18E2 (Reilly, Deitz, et al., in prep). We identified homologous proteins using a reciprocal best exonerate approach [53]: *D. melanogaster* proteins were used to align and extract proteins from each genome reference sequence and the resultant proteins were aligned back to the *D. melanogaster* genome to verify the same protein was returned. The longest *D. melanogaster* transcript was selected for each gene and PRANK was used to create multispecies CDS alignments using the “codon” setting [54]. This resulted in 13,169 CDS alignments, of which 11,496 (87%) passed the filtering criteria and were used in the clustering analysis. From this alignment, we quantified clustering in *D. melanogaster* and *D. simulans* using *D. yakuba* as the outgroup.

To insure high-quality alignments, several filtering criteria were imposed on each CDS alignment. 1) The length of the alignment must be a multiple of three and must not contain premature stop codons. 2) Any gaps in the alignments must be a multiple of 3 nucleotides in length, so frameshifts are not introduced. 3) Amino acid substitutions must comprise less than 20% of the protein, as rates higher than this likely indicate an incorrect alignment. 4) Gaps must comprise less than 20% of the alignment length, as a proxy for genome assembly quality.

### Clustering calculations

For each protein alignment, we identified codons containing either a synonymous (DS) or non-synonymous substitution (DN) between each pair of species. Each codon is allowed to contain only one substitution and we do not attempt to identify the order in which the substitutions occurred. In other words, a codon may have two or three nucleotide substitutions, but it is only considered a single DN or DS. To calculate DNDN and DSDS clustering, for each protein, we take all pairs of non-synonymous substitutions or synonymous substitutions, respectively, and calculate the distance, in codons, between each pair. After calculating the distances between all pairs of DNDN and DSDS for each protein, genome-wide, we count the number of pairs separated by *x* codons, for *x* ∈ [1,500], to get a non-normalized clustering score for each *x*. This clustering score for DNDN and DSDS of each protein is normalized by the analytical expectation for all pairs of distances between uniformly distributed substitutions as

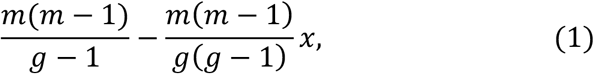

where *m* is the number of codon substitutions (either DN or DS) and *g* is the protein length in codons.

This equation was verified by simulation of substitutions sampled from a uniform distribution, without replacement. This clustering metric was further normalized by the average clustering score over an 80-120 codon window. The correlation between distance in the primary sequence and distance between alpha carbons in the 3D folded proteins breaks down well before an 80 codon length scale (Figure 2), thus we do not expect to observe clustering at this length scale. Furthermore, the clustering distribution asymptotes at this length scale, matching our hypothesis. In all plots, data was smoothed with a sliding-window average of 5 codons. This clustering metric is an aggregated, genome-wide metric which makes it difficult to identify specific genes with above-average clustering scores. We calculate significance of DNDN over the first 20 codons using a chi-square goodness of fit test where our expectation is based either on the null distribution of uniformly distributed mutation (described in equation 1) or DSDS, as this is meant to be a control.

### Quantification of clustering using protein structures

To check that distance in codons along the primary sequence is a good proxy for distance in the folded protein, we calculated clustering with distance in 3D space as our distance metric for *Saccharomyces* and primates. We downloaded all the PDB files associated to *S. cerevisiae* and *H. sapiens* from www.rcsb.org. PDB files were parsed using the PDB module of Biopython [55]. CDS alignments were matched to their corresponding 3D structure information based on each gene’s associated PDB ID and Chain ID. The sequence within the PDB file was verified against the CDS alignment sequence. In cases where there were multiple PDB files for a given gene, the PDB file containing the longest sequence was used and when there were multiple of these, one was chosen arbitrarily. This resulted in a total of 1,628 genes (38%) and 3,479 genes (37%) that we were able to use to quantify 3D clustering in *Saccharomyces* and primates, respectively. Clustering was repeated as before, first by identifying synonymous and non-synonymous sites, then calculating the distance in angstroms between each pair of synonymous sites and non-synonymous sites. The number of DNDN and DSDS in bins of 2Å were counted. The normalization for 3D DNDN and DSDS was calculated by counting the total number of all pairs of amino acid residues in 2Å bins and normalizing this to the observed number of DNDN or DSDS.

### Lineage-specific clustering

To break the dependence of substitutions on one another, we can polarize substitutions into the lineage in which they arose and then calculate clustering between lineages. This will measure clustering due to lack of constraint since substitutions between species cannot interact with one another. Thus, the within-lineage clustering in excess of between-lineage clustering should be due exclusively to interactions between substitutions. Following previous work [30], we used an outgroup to polarize substitutions into the lineage in which they arose using parsimony (such that the number of substitutions that occurred ancestrally was minimized). Substitutions that could not be polarized were omitted from further analysis. DNDN was calculated within and between species pairs. To calculate within species DNDN, we take all pairs of non-synonymous substitution within one species and count the number of pairs separated by each distance, in codons, for *x* ∈ [1,500]. Again, this distribution is normalized by the null distribution of uniformly distributed substitutions, as given in equation 1. To calculate between species DNDN, we take all pairs of non-synonymous substitutions such that the two substitutions are from different species and count the number of pairs at each distance for *x* ∈ [1,500]. We then normalize this by the analytical distribution of the pairs of two different “types” of substitutions, where each type is uniformly distributed as

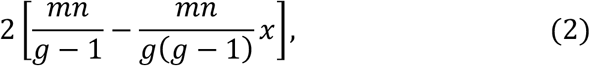

where *m* is the number of DN in species one, *n* is the number of DN in species two, and *g* is the protein length. We are mainly interested in the within species clustering significantly in excess of between species clustering. Thus, we calculate the significance of within species clustering over the first 20 codons using a chi-square goodness of fit test using between species clustering as our expectation.

Since we observe an excess of within-lineage clustering, we wanted to examine whether certain classes of amino acid substitutions were over represented in the non-synonymous clustering we previously observed. To that end, we examined three properties of amino acids, size, polarity, and charge, and classify each non-synonymous substitution as increasing or decreasing the previously mentioned properties. Then, we can classify each pair of non-synonymous substitutions as compensatory or reinforcing. We represent compensatory and reinforcing pairs of substitutions as the fraction of DNDN in which the change in the substituted amino acid are anti-correlated (compensatory) or correlated (reinforcing). To calculate the significance of within species compensatory clustering, we use a chi-square contingency table to compare the frequency of compensatory pairs within species to the frequency of compensatory pairs between species over the first 10 codons. The same was done for reinforcing clustering. We used 10 codons as the length-scale of clustering appeared to be shorter.

### Simulations of protein-folding stability

To address the extent to which purifying selection plays a role in the clustering signal, we conducted simulations where we maintained the folding energy of the protein (as a proxy for protein function/fitness) to explore whether we could recapitulate clustering due to epistasis under a model of exclusively purifying selection. FoldX, a forcefield-based protein simulation software package, was used to calculate the folding energy of proteins with substitutions, as it has been previously used to simulate epistasis [25].

Previous studies show that FoldX is more accurate when predicting folding energy for small proteins so we chose to simulate substitutions in three, short, monomeric proteins: Lysine-, arginine-, ornithine-binding protein from *Salmonella typhimurium*; NmrA-like family domain containing protein 1 from *Homo sapiens*; and Mini spindles TOG3 from *D. melanogaster*, corresponding to PDB IDs: 1LAF, 2WM3, and 4Y5J, respectively. Additionally, these proteins had no missing residues in their PDB files, low clashscores, and few sidechain outliers, making them well-suited for FoldX simulation. Lastly, these proteins have broadly differing geometries: 1LAF is globular while 4Y5J has a more linear structure and 2WM3 is intermediate between the two. This can be seen from the plots of distance measured in codons (along the primary sequence) vs distance in 3D folded space (Figure S3).

Protein structures were downloaded from www.rcsb.org using the aforementioned PDB IDs. We followed the methodology from Shah et al. [25]; in brief, the protein structure was preprocessed with RepairPDB, and BuildModel was used to estimate the change in the folding energy of the protein as a result of the substitution we simulated.

To simulate substitutions, we started from the wild-type protein sequence, generated 10 single amino acid mutations (with the restriction that the new mutations could not revert the previous), and calculated the change in free folding energy of each mutant (ΔΔG). We then calculated a probability of fixation for each mutant using a Gaussian fitness function,

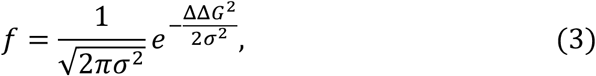

and a standard Moran process,

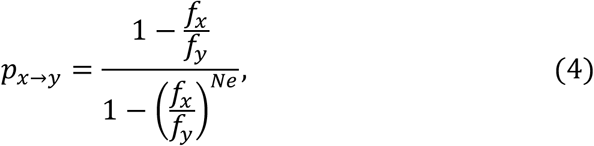

where *σ*^2^ = 1450 and *Ne* = 10,000 [25]. The probability of fixation,

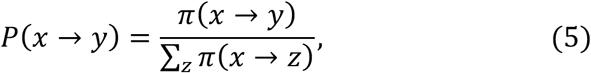

was scaled to 1. At each step, the mutant is selected randomly, proportional to its fixation probability. This process is repeated for the desired number of substitutions (two). We additionally conducted simulations using a piecewise fitness function, in which decreased stability is given a fitness penalty, while increased stability has equivalent stability to wild type.

We calculated clustering for the simulations as we did in the real data, using equation 1. We additionally wanted to calculate clustering due solely to epistasis. That is to say, controlling for clustering due to reduced constraint along the protein sequence. We calculated the null distribution of clustering due to constraint by resampling the first and second substitutions separately to remove any epistatic interactions between the pair and used this to normalize the mutation pair clustering. Finally, to enrich for pairs of substitutions with increased levels of epistatic interactions, we ordered pairs based on the difference between the additive ΔΔG of the single mutants vs the double mutant: |ΔΔ*G*_1_ + ΔΔ*G*_2_ – ΔΔ*G*_1,2_| and selected the top *n*% of them.

## Acknowledgements

Thanks to Premal Shah for sharing code for FoldX simulations. Thanks to Clair Han and Patrick Reilly for scientific discussions. This work was funded by R01 GM115523 to P.A.

