## Supplement for "Epistasis and physico-chemical constraints contribute to spatial clustering of amino acid substitutions in protein evolution"

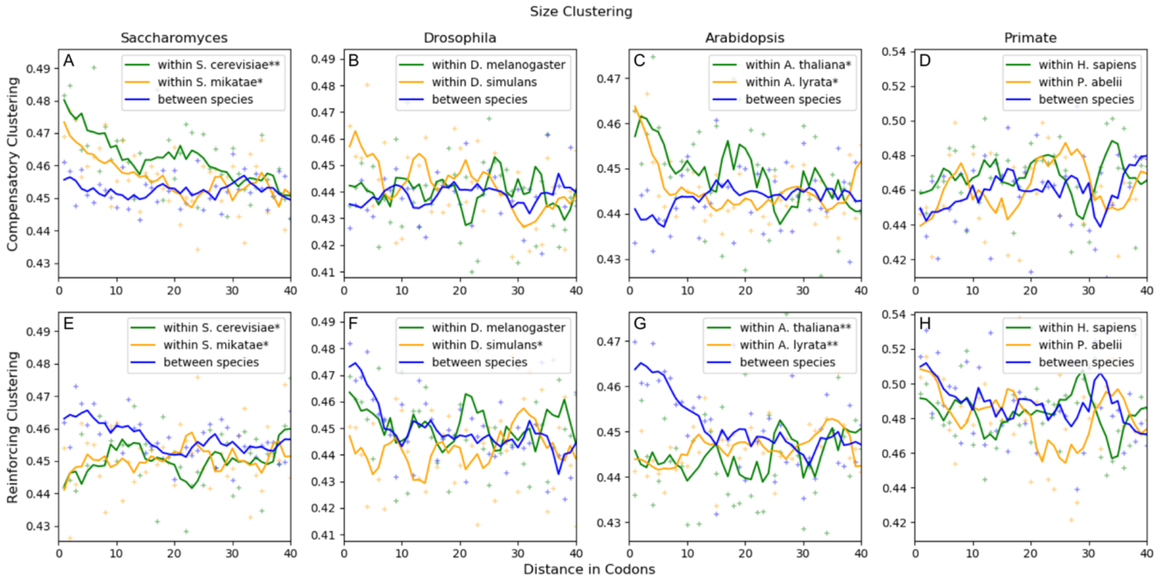


**Supplementary figure 1:** Pairs of non-synonymous substitutions causing size differences typically exhibit similar patterns of compensatory clustering within species and reinforcing clustering between species as seen for charge clustering (Figure 4). The top panels (A, B, C, D) show the fraction of non-synonymous substitution pairs exhibiting size compensating interactions. The bottom panels (E, F, G, H) show the fraction of non-synonymous substitution pairs exhibiting reinforcing interactions. Green and orange indicates pairs that occur within species and blue indicates pairs that occur between species. P-values are indicated in the legend by: ** $p < 10^{-5}$; * $10^{-5} < p < 0.05$. See Table S4 for p-values .The following comparisons are shown: A, E) *S. cerevisiae* and *S. mikatae*, outgroup: *S. kudriavzevii*. B, F) *D. melanogaster* and *D. simulans*, outgroup: *D. yakuba*. C, G) *A. thaliana* and *A. lyrata*, outgroup: *C. rubella*. D, H) *H. sapiens* and *P. pygmaeus abelii*, outgroup: *P. anubis*.

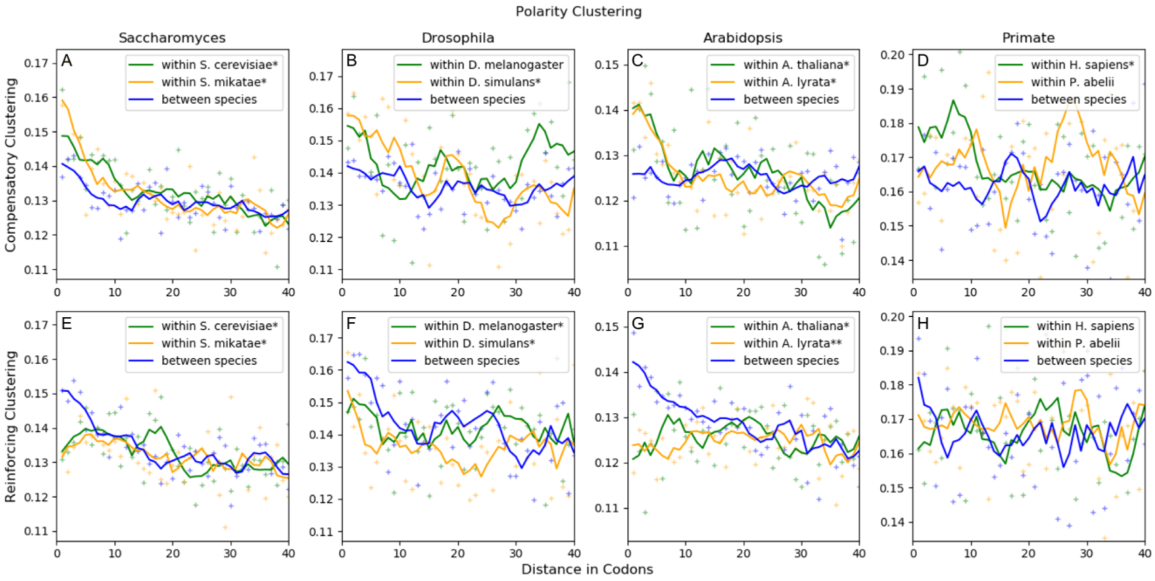


**Supplementary figure 2:** Pairs of non-synonymous substitutions causing polarity differences typically exhibit similar patterns of compensatory clustering within species and reinforcing clustering between species as seen for charge clustering (Figure 4). The top panels (A, B, C, D) show the fraction of non-synonymous substitution pairs showing polarity compensation, while the bottom panels (E, F, G, H) show the fraction of non-synonymous substitution pairs showing reinforcing polarity changes. As before, within species pairs are shown in green and orange, while between species pairs are shown in blue. P-values are indicated in the legend by: ** $p < 10^{-5}$; * $10^{-5} < p < 0.05$. See Table S5 for p-values. The following comparisons are shown: A, E) *S. cerevisiae* and *S. mikatae*, outgroup: *S. kudriavzevii*. B, F) *D. melanogaster* and *D. simulans*, outgroup: *D. yakuba*. C, G) *A. thaliana* and *A. lyrata*, outgroup: *C. rubella*. D, H) *H. sapiens* and *P. pygmaeus abelii*, outgroup: *P. anubis*.

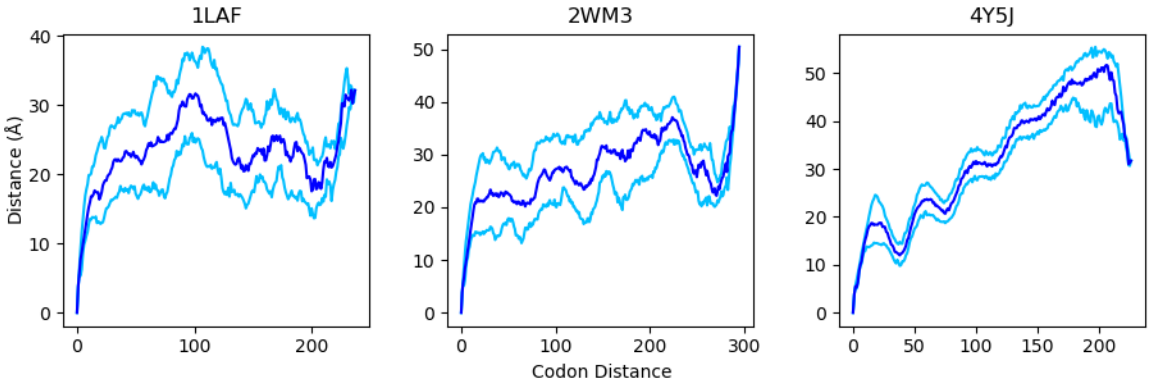


**Supplementary figure 3:** Distance in 3D space (in Å) vs distance along the primary sequence (in codons) for the three proteins we simulated. Median distance (blue) and 1st and 3rd quartiles (light blue) are shown. Protein 1LAF is relatively globular, as the 3D distance rapidly asymptotes, while protein 4Y5J maintains the linear trend throughout most of the protein.

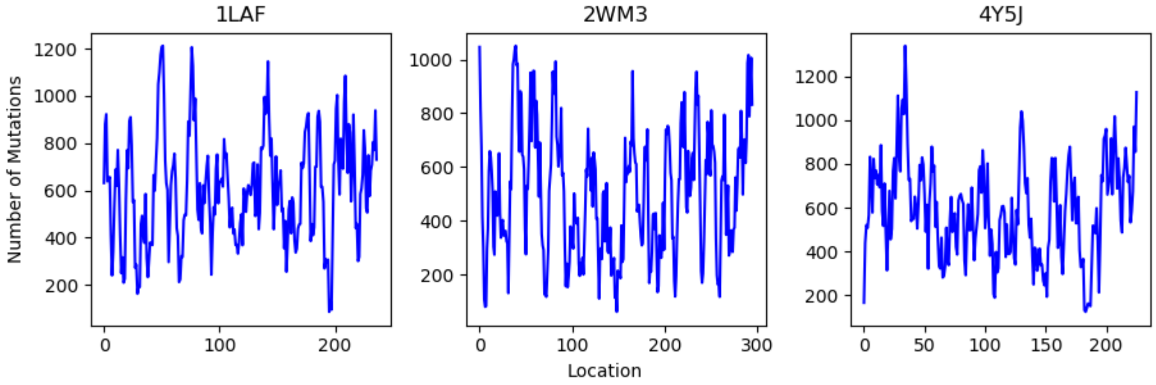


**Supplementary figure 4:** Distributions of simulated mutations across the three proteins. While it does appear that there is substantial variability in the locations of the simulated mutations, there are no obvious patterns indicating strong local constraint in any specific region of the proteins.

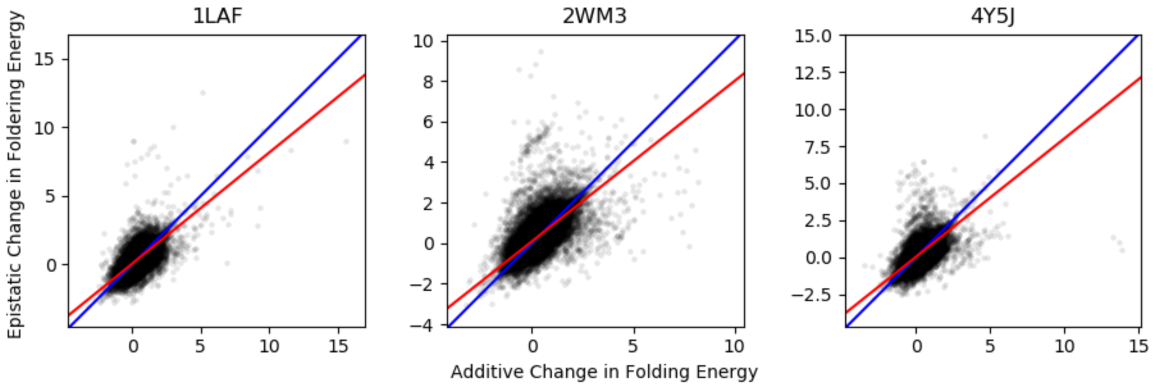


**Supplementary figure 5:** Epistatic change in folding energy ($\Delta\Delta G_{1,2}$) vs additive change in folding energy ($\Delta\Delta G_1 + \Delta\Delta G_2$) for all three proteins simulated. As in Figure 5A, the epistatic change in folding energy is less than the additive prediction for all three proteins.

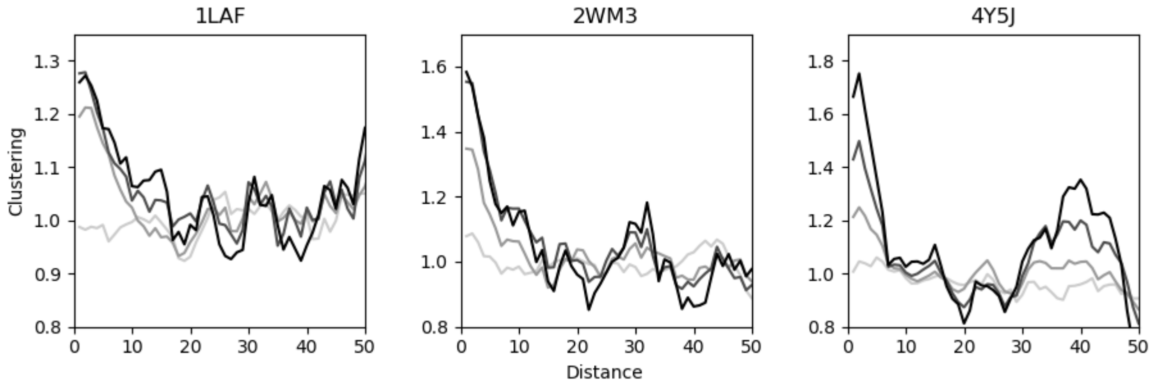


**Supplementary figure 6:** Clustering of all pairs of simulated mutations (lightest gray) shows no clustering in any of the three proteins we simulated. However, we do observe clustering when taking the top 50%, 20% and 10% of the most epistatic pairs of mutations (increasing darkness, as in Figure 5).

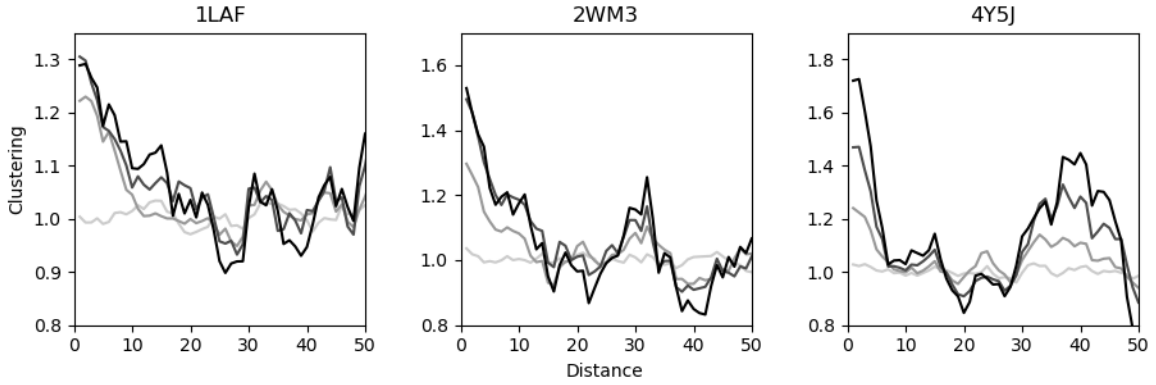


**Supplementary figure 7:** Clustering after normalizing for local variation in constraint across the protein. Qualitatively, the patterns of clustering are the same as the ones seen in Figure S6, indicating that variation in constraint across the proteins is not driving the patterns of clustering. Again, epistasis thresholds are shown in increasingly dark gray over the top 100%, 50%, 20%, and 10% of the most epistatic pairs of mutations.

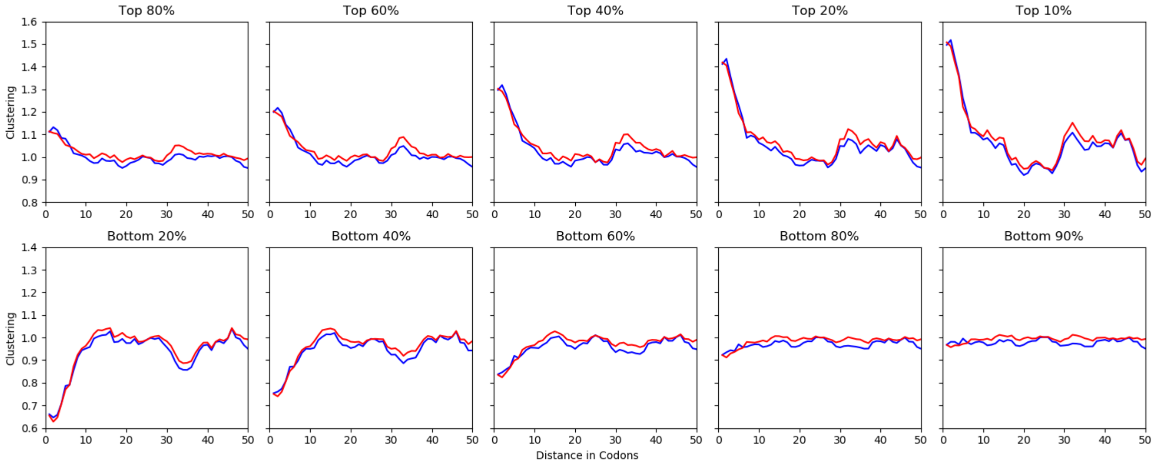


**Supplementary figure 8:** Clustering for the aggregated mutations across all three proteins at different epistasis thresholds. The top row shows the clustering of the top n% epistatic pairs of mutations, while the bottom row shows the complement (i.e. the bottom (1-n)% least epistatic pairs). With increasingly stringent epistasis thresholds, the clustering gets more severe. Interestingly, taking even a modest subset of the pairs of mutations (i.e. “Top 80%”), we begin to see patterns of clustering. This is explained by the fact that almost all of the pairs of mutations being removed occur at large distances (“Bottom 20%” – the most additive 20% of simulated mutation pairs).

| DNDN | *Saccharomyces* | *Drosophila* | *Arabidopsis* | Primates |
|---|---|---|---|---|
| vs uniform | $< 10^{-300}$ | $< 10^{-300}$ | $< 10^{-300}$ | $1.1 \times 10^{-109}$ |
| vs DSDS | $< 10^{-300}$ | $< 10^{-300}$ | $< 10^{-300}$ | $2.6 \times 10^{-78}$ |

**Table S1:** Non-polarized clustering p-values. P-values comparing the distribution of DNDN to the expectation of uniformly distributed substitutions (top row), or to the DSDS distribution (bottom row). All p-values are highly significant.

| | *Saccharomyces* | *Drosophila* | *Arabidopsis* | Primates |
|---|---|---|---|---|
| Within species 1 | $7.3 \times 10^{-11}$ | $2.0 \times 10^{-11}$ | $1.9 \times 10^{-13}$ | 0.010 |
| Within species 2 | $7.9 \times 10^{-7}$ | $5.3 \times 10^{-10}$ | $2.8 \times 10^{-12}$ | $4.1 \times 10^{-42}$ |

**Table S2:** Polarized clustering p-values. P-values comparing the distribution of within species clustering to that of the between species clustering. In the table, species 1 corresponds to *S. cerevisiae*, *D. melanogaster*, *A. thaliana*, and *H. sapiens*, while species 2 corresponds to *S. mikatae*, *D. simulans*, *A. lyrata*, and *P. pygmaeus abelii*.

| | | *Saccharomyces* | *Drosophila* | *Arabidopsis* | Primates |
|---|---|---|---|---|---|
| Compensatory | Within species 1 | $1.4 \times 10^{-16}$ | $3.9 \times 10^{-10}$ | $2.1 \times 10^{-19}$ | 0.076 |
| | Within species 2 | $4.1 \times 10^{-10}$ | $5.6 \times 10^{-13}$ | $1.0 \times 10^{-19}$ | $2.8 \times 10^{-4}$ |
| Reinforcing | Within species 1 | $1.0 \times 10^{-15}$ | $1.7 \times 10^{-11}$ | $2.2 \times 10^{-9}$ | 0.12 |
| | Within species 2 | $2.7 \times 10^{-10}$ | $1.8 \times 10^{-8}$ | $1.6 \times 10^{-26}$ | 0.26 |

**Table S3:** Charge clustering p-values. P-values comparing the ratio of within species DNDN to between species DNDN clustering to the ratio of within species charge compensating or reinforcing DNDN to between species charge compensating or reinforcing DNDN. In the table, species 1 corresponds to *S. cerevisiae*, *D. melanogaster*, *A. thaliana*, and *H. sapiens*, while species 2 corresponds to *S. mikatae*, *D. simulans*, *A. lyrata*, and *P. pygmaeus abelii*.

| | | *Saccharomyces* | *Drosophila* | *Arabidopsis* | Primates |
|---|---|---|---|---|---|
| Compensatory | Within species 1 | $2.5 \times 10^{-7}$ | 0.78 | $9.5 \times 10^{-5}$ | 0.13 |
| | Within species 2 | $2.1 \times 10^{-3}$ | 0.053 | $3.2 \times 10^{-4}$ | 0.41 |
| Reinforcing | Within species 1 | $5.0 \times 10^{-5}$ | 0.32 | $7.1 \times 10^{-7}$ | 0.27 |
| | Within species 2 | $3.4 \times 10^{-4}$ | $1.6 \times 10^{-3}$ | $1.1 \times 10^{-9}$ | 0.46 |

**Table S4:** Size clustering p-values. P-values comparing the ratio of within species DNDN to between species DNDN clustering to the ratio of within species size compensating or reinforcing DNDN to between species size compensating or reinforcing DNDN. In the table, species 1 corresponds to *S. cerevisiae*, *D. melanogaster*, *A. thaliana*, and *H. sapiens*, while species 2 corresponds to *S. mikatae*, *D. simulans*, *A. lyrata*, and *P. pygmaeus abelii*.

| | | *Saccharomyces* | *Drosophila* | *Arabidopsis* | Primates |
|---|---|---|---|---|---|
| Compensatory | Within species 1 | $2.6 \times 10^{-4}$ | 0.29 | $3.5 \times 10^{-3}$ | 0.012 |
| | Within species 2 | $1.3 \times 10^{-4}$ | $3.7 \times 10^{-3}$ | $2.0 \times 10^{-5}$ | 0.47 |
| Reinforcing | Within species 1 | 0.038 | 0.031 | $4.2 \times 10^{-5}$ | 0.76 |
| | Within species 2 | $1.4 \times 10^{-3}$ | $9.3 \times 10^{-3}$ | $8.7 \times 10^{-12}$ | 0.79 |

**Table S5:** Polarity clustering p-values. P-values comparing the ratio of within species DNDN to between species DNDN clustering to the ratio of within species polarity compensating or reinforcing DNDN to between species polarity compensating or reinforcing DNDN. In the table, species 1 corresponds to *S. cerevisiae*, *D. melanogaster*, *A. thaliana*, and *H. sapiens*, while species 2 corresponds to *S. mikatae*, *D. simulans*, *A. lyrata*, and *P. pygmaeus abelii*.
